# The relationship between resting-state functional connectivity, antidepressant discontinuation and depression relapse

**DOI:** 10.1101/2020.02.06.937268

**Authors:** Isabel M. Berwian, Julia G. Wenzel, Leonie Kuehn, Inga Schnuerer, Lars Kasper, Ilya M. Veer, Erich Seifritz, Klaas E. Stephan, Henrik Walter, Quentin J. M. Huys

**Author notes:** Equal contribution (Henrik Walter, Quentin J. M. Huys). **Corresponding author**: Isabel Berwian.

## Abstract

The risk of relapsing into depression after stopping antidepressants is high, but no established predictors exist. Resting-state functional magnetic resonance imaging (rsfMRI) measures may help predict relapse and identify the mechanisms by which relapses occur. rsfMRI data were acquired from healthy controls and from patients with remitted major depressive disorder on antidepressants. Patients were assessed a second time either before or after discontinuation of the antidepressant, and followed up for six months to assess relapse. A seed-based functional connectivity analysis was conducted focusing on the left subgenual anterior cingulate cortex and left posterior cingulate cortex. Seeds in the amygdala and dorsolateral prefrontal cortex were explored. 44 healthy controls (age: 33.8 (10.5), 73% female) and 84 patients (age: 34.23 (10.8), 80% female) were included in the analysis. 29 patients went on to relapse and 38 remained well. The seed-based analysis showed that discontinuation resulted in an increased functional connectivity between the right dorsolateral prefrontal cortex and the parietal cortex in non-relapsers. In an exploratory analysis, this functional connectivity predicted relapse risk with a balanced accuracy of 0.86. Further seed-based analyses, however, failed to reveal differences in functional connectivity between patients and controls, between relapsers and non-relapsers before discontinuation and changes due to discontinuation independent of relapse. In conclusion, changes in the connectivity between the dorsolateral prefrontal cortex and the posterior default mode network were associated with and predictive of relapse after open-label antidepressant discontinuation. This finding requires replication in a larger dataset.

## 1 Introduction

A subset of those suffering from Major Depressive Disorder (MDD) achieve remission with antidepressant medication (ADM)(1). However, this does not imply that the illness is cured as one in three of those achieving remission experience a relapse within six months after ADM discontinuation(2). Indeed, the burden of depressive illness is in no small part due to their chronic nature with frequent relapses over the lifetime(3; 4). Hence, the management of relapses is of paramount importance. In this situation, predictors of relapse risk are urgently needed to guide decision-making at key decision points such as the discontinuation of ADM after remission has been achieved.

Unfortunately, standard demographic and clinical variables appear to have very limited predictive power(5), requiring attention to be turned to more complex measurements. Here, we examined resting-state functional connectivity - a complex yet clinically feasible neuroimaging procedure(6).

Indeed, a substantial body of research has examined resting-state functional connectivity (RSFC) in the context of depression. Abnormal RSFC in MDD has been observed in three core networks, i.e. the so-called “default mode network” (DMN), the central executive network (CEN) and the salience network (SN). Specifically, in MDD, the connectivity within the anterior and posterior DMN is increased and connectivity between the anterior DMN and the affective and salience network is also altered(7; 8). However, a reduction in connectivity within the DMN has also been reported amongst medicated patients and those with recurrent illness(9). Connectivity within the CEN and SN and connectivity to the posterior DMN are reduced(7; 8).

In addition to abnormal RSFC in current depression(10; 11; 12), the subgenual anterior cingulate cortex (sgACC) appears to have a particularly central role in relation to treatment response, remitted depression and relapse. First, neuroimaging measures of metabolic activity within this region have been shown to have predictive power for response to antidepressant and other treatments(13; 14). Second, connectivity changes in these regions have also been observed in remitted patients off medication and in populations with correlates of depression such as trauma, childhood maltreatment, subclinical depression and familial risk in children. These include increased connectivity between the sgACC and both the medial prefrontal cortex (mPFC) and the posterior cingulate cortex (PCC), as well as bidirectional changes in connectivity between the sgACC and the dorsolateral PFC (dlPFC), the amygdala and the hippocampus((15; 16; 17; 18; 19; 20; 21; 22; 23; 24); though see also (25)). Third, and most importantly, decreased interhemispheric sgACC connectivity from the left to the right has been suggested as a marker of resilience to relapse in patients off medication(26), while increased connectivity between the sgACC and regions in the CEN differentiates patients who will experience a relapse and those who will not(27).

The results of studies examining the impact of ADM initiation on connectivity are heterogeneous(28; 29; 30; 31; 32; 33; 34; 35; 36; 37; 38; 39). One of the more replicated findings is a decrease in connectivity between PCC and various brain regions (including amygdala(31), right inferior temporal gyrus(36), right inferior frontal gyrus(33)), suggesting a normalisation of increased PCC connectivity with these regions in comparison to the prior, medication-free, state. Increases between the PCC and the ACC and mPFC have been reported in patients who receive treatment, but do not remit(32).

Outstanding gaps in the literature include characterization of RSFC in the remitted medicated state, difference in RSFC between patients who relapse and remain well after antidepressant discontinuation and the neurobiological effects underpinning discontinuation. We aimed to address these gaps in the context of the AIDA study - a two-centre observational randomized study that followed patients as they discontinued their ADM for six months. The fundamental goal of the study was to identify biomarkers of relapse risk and uncover neurobiological changes due to discontinuation. To this end, we examined 1) whether RSFC connectivity differs between remitted medicated patients and controls; 2) whether pre-discontinuation RSFC differs between patients who relapse after discontinuation and those who remain well; 3) the effects of discontinuation on RSFC; and 4) to what extent changes in RSFC due to discontinuation are related to subsequent relapses. Based on the reviewed literature, we focused on the sgACC as seed region to address questions 1 and 2 and on the PCC for questions 3 and 4.

## 2 Methods and Materials

### 2.1 Study Design

The AIDA study design is depicted in Figure 1. Briefly, we focused on remitted patients on ADM who wanted to discontinue their medication. Participants were randomised to one of the two study arms. In group MA1-D-MA2, participants underwent the first main assessment (MA1), then gradually discontinued their medication over up to 18 weeks, and then underwent a second main assessment (MA2). In group MA1-MA2-D, participants underwent two main assessments and then discontinued. Hence, the group name indicates the order of events and “D” indicates discontinuation.

**Figure 1.**
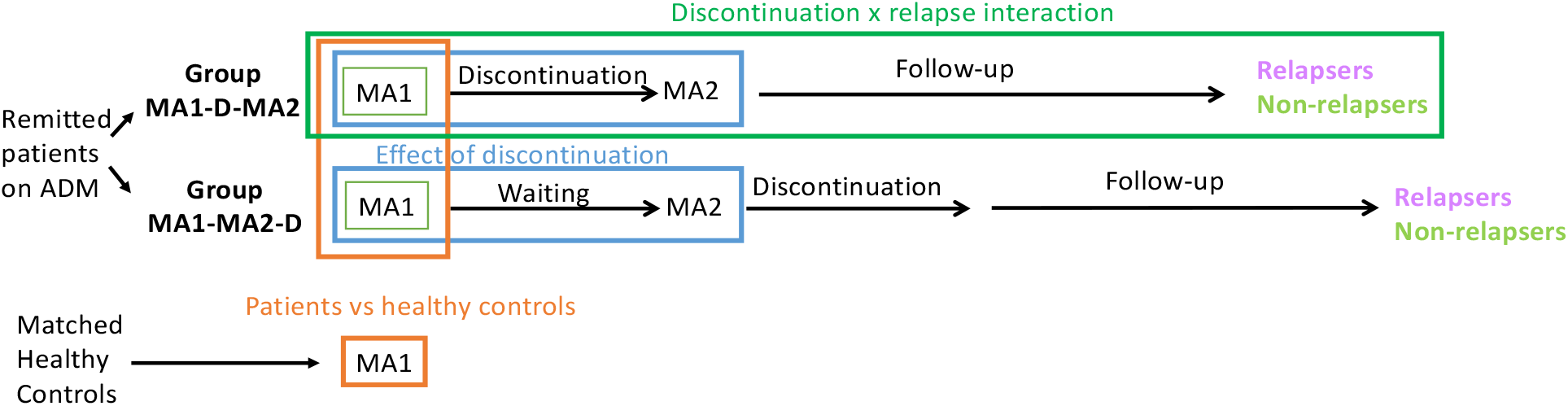
Study Design: Remitted patients on antidepressant medication (ADM) and matched healthy controls were included in the study and assessed during main assessment 1 (MA1). Patients were randomised to groups MA1-D-MA2 or MA1-MA2-D, where the name indicates the order of events and “D” indicates discontinuation. In group MA1-D-MA2, they underwent MA1, then discontinued their medication followed by main assessment 2 (MA2). In group MA1-MA2-D, they underwent MA1 and MA2 first and only discontinued thereafter. After discontinuation, all patients were followed up for six months to ascertain relapses. Comparison of patient and control groups at MA1 was used to examine the remitted medicated state cross-sectionally (orange). Data collected at MA1 from all patients who subsequently completed the study as relapsers or non-relapsers, i.e. relapsers and non-relapsers from both groups combined were examined to examine associations with relapse. The interaction between time point (MA1 and MA2) with group (MA1-D-MA2 and MA1-MA2-D) examined the impact of discontinuation (blue). Finally, the interaction between time point (MA1 and MA2) and relapse in group MA1-D-MA2 was used to examine changes due to discontinuation that led to relapse or resilience (dark green).

After discontinuation, all patients were contacted by telephone at weeks 1, 2, 4, 6, 8, 12, 16 and 21 to assess relapse status. If subjects exhibited signs of a relapse, a in-person structured clinical interview was performed (SCID-I(40)) to establish the presence of a DSM-IV-TR major depressive episode. If this was present, participants underwent a final assessment (FA) at that point, otherwise the final assessment took place in week 26. Healthy controls (HC) matched for age, sex and education were only assessed once (MA1).

All patients underwent a resting-state functional magnetic resonance imaging (rsfMRI), self- and observer-rated reports during each main assessment (MA1 and MA2) and HC once at MA1. See supplementary section S1.2 for observer-rated and self-report measures. Participants were recruited between July 2015 and January 2018.

### 2.2 Participants

The AIDA study recruited participants who had experienced one severe(41) or multiple depressive episodes; had initiated antidepressant treatment during the last depressive episode and now achieved stable remission; and had reached the decision to discontinue their medication independently from and prior to study participation. Detailed inclusion and exclusion criteria are listed in the supplementary section S1.1. All participants gave informed written consent and received monetary compensation for the time of participation. Ethical approval for the study was obtained from the cantonal ethics commission Zurich (BASEC: PB_2016-0.01032; KEK-ZH: 2014-0355) and the ethics commission at the Campus Charité-Mitte (EA 1/142/14). All experiments were performed in accordance with relevant guidelines and regulations.

### 2.3 Seed Region Selection

Seven seeds of interest (Figure 2A) were identified based on existing literature. The sgACC seeds with MNI coordinates +/−4, 21, −8 were chosen as they had been used in several previous studies with patients with remitted MDD(15; 19; 20; 21; 27) and clearly mapped on the anatomical region of the sgACC during visual inspection. For the left PCC we chose the MNI coordinates −3, −39, 39 as identified by Fox and colleagues(42) as a central part of the task-negative network and were reported to have normalised RSFC after ADM initiation(35). dlPFC coordinates (MNI +/−37, 26, 31) were taken from Sheline and colleagues(43) and have been used widely in the literature. Unless stated otherwise, seeds were created by putting a 6mm sphere around the coordinates. Amygdala seeds were extracted from the Harvard-Oxford Subcortical Probability Atlas by including all voxels with at least 80% probability of being in the amygdala. For each analysis, one seed was designated a priori as the main seed of interest and the other six as exploratory seeds.

**Figure 2.**
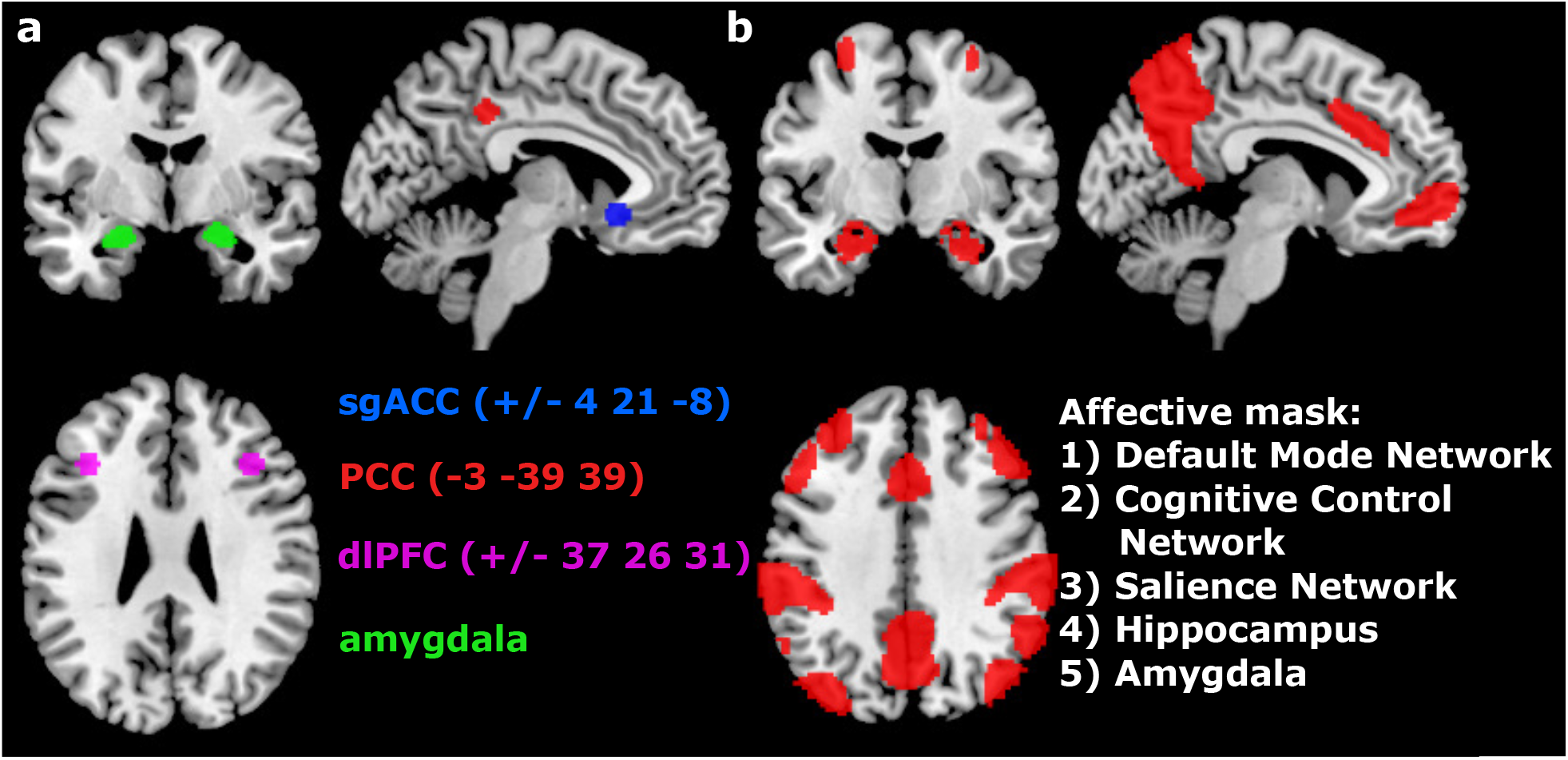
Seeds and Mask: Depicted are the seeds (A) used in the analysis and the affective mask (B) containing all voxels of interest. sgACC = subgenual anterior cingulate cortex; PCC = posterior cingulate cortex, dlPFC = dorsolateral prefrontal cortex

### 2.4 Resting-state fMRI analysis

#### 2.4.1 Image acquisition and preprocessing

The sequences used for image acquisition are described in the supplementary section S1.4. Imaging data was preprocessed with a FSL (FMRIB Software Library v5.0) pipeline detailed in supplementary section S1.5.

#### 2.4.2 First-level analyses

First level analyses were conducted with the CONN toolbox(44). We included motion regressors (see section motion correction for details), 10 regressors computed using aCompCor in the CONN toolbox (5 principal components of white matter and cerebrospinal fluid signals each), and bandpass filtered the data to retain frequencies between 0.01 and 0.1 Hz. All covariates were also regressed out of the seed regions. Then, the average signal of all voxels within subject-specific grey matter masks of each seed was computed and correlated with all voxels in the brain. Resulting correlation coefficients were z-transformed for second-level analyses.

Analyses to control for effects of motion and study site are described in detail in supplementary section S1.6 and S1.7. Subjects with a framewise displacement (FD) of 1mm from one volume to the next and with a mean SNR smaller than 30 were excluded from further analyses.

#### 2.4.3 Second-level analyses

Second-level analyses were conducted with SPM12 (version: v7219). For all analyses with our chosen main seed, we report peak-level results family-wise error (FWE) corrected at the voxel level at 0.05 and depict the respective clusters in figures. Altered RSFC of the selected seeds in acute and remitted depression has mainly been reported for voxels which are part of the DMN, the CEN, the SN, the amygdala and the hippocampus(8; 7; 15; 17; 19; 20; 21; 24). As FWE-corrected results depend on the search volume of the analysis, we constructed an “affective mask” containing all voxels within the DMN, the CEN and the SN, as well as voxels within anatomical masks of the bilateral amygdalae and the hippocampi (Figure 2B) as these areas show abnormal RSFC in affective disorders. Details on included areas in the mask can be found in supplementary section S1.8 Exploratory analyses were considered significant after Bonferroni correction for all six seeds at p=0.0083.

Our first analyses focused on the first assessment MA1. To identify abnormalities persisting into the remitted medicated state we compared RSFC between the left sgACC and the affective mask between patients and HC using an independent-sample t-test. To identify markers of prospective relapse, we repeated this analysis comparing patients who did and did not go on to relapse. Exploratory analyses examined the RSFC between the affective mask and the right sgACC, the left PCC, right and left amygdala and right and left dlPFC seed regions.

Next, we examined the impact of discontinuation. To test the hypothesis that RSFC between the PCC and the affective mask would increase due to discontinuation, we applied a mixed-design analysis of variance (ANOVA) with group (MA1-D-MA2 and MA1-MA2-D) as between-subject factor and time point as within-subject factor (MA1 and MA2). Where significant interaction effects were found, we conducted the following post-hoc tests: paired sample t-test in group MA1-D-MA2 to identify changes related to discontinuation; paired sample t-test in group MA1-MA2-D to investigate test-retest reliability and an independent sample t-test between groups MA1-D-MA2 and MA1-MA2-D at MA1 to verify that no random allocation differences occurred prior to discontinuation. We repeated this analyses using the right and left sgACC, right and left amygdala and right and left dlPFC as seeds for exploratory analyses.

Finally, we attempted to gain insight into how the effects of discontinuation might relate to relapse by examining the interaction of discontinuation and relapse. For this, we only included patients from group MA1-D-MA2 and applied a mixed-design ANOVA with time point as within-subject factor (MA1 and MA2) and relapse status (relapsers and non-relapsers) as between subject factor. Post-hoc paired sample t-tests were performed within the relapse group to identify changes related to relapse and paired t-tests in the no-relapse group to identify changes related to resilience.

Additional sanity checks and exploratory analyses are described in supplementary section S1.9.

#### 2.4.4 Covariates

RSFC that were found to differ significantly between or within groups were correlated with several covariates including age, gender and site as covariates of no interest and rumination score, residual depression score, medication load and disease severity. Computation of medication load and severity factor is outlined in supplementary section S1.3.

#### 2.4.5 Post-hoc prediction analyses

As suggested by a reviewer we examined if associations between RSFC and relapse would also predict relapse risk out-of-sample. To this end, we fitted logistic regressions nested in a leave-one-out cross-validation for the main significant cluster in the interaction analyses that indicated differences in changes in RSFC between MA1 und MA2 in relapsers and non-relapsers. Group membership of the left-out participant was predicted using parameter estimates (regression weights) obtained from the participants included in the fit. Finally, the balanced accuracy for predicted group memberships was computed.

## 3 Results

### 3.1 Participants

The consort diagrams (Figure S1 and Figure S2) show numbers of patients and controls at each study stage and reasons for dropouts. After exclusion, 84 patients (mean [SD] age, 34.23 (10.8) years, 17 men [20%]) and 44 controls (mean [SD] age, 33.8 (10.5) years, 12 men [27%]) were included at MA1. 61 (73%) patients and 27 (61%) controls were assessed in Zurich, the remaining participants in Berlin. Further details on demographic and clinical characteristics are displayed in Table 1. Patients scored higher on the rumination brooding subscale. This was particularly accentuated in patients who went on to relapse. 69% of our sample took a selective serotonin reuptake inhibitor, 27% a serotonin-norepinephrine reuptake inhibtor and 4% an antidepressant from a different class.

**Table 1:** Participant characteristics

| Variable<sup>a</sup> | Patients vs. Healthy controls |  |  | Relapsers vs. Non-relapsers |  |  |
| --- | --- | --- | --- | --- | --- | --- |
|  | Patients (n = 84) | Healthy controls (n = 44) | P value | Relapsers (n = 29) | Non-relapsers (n = 38) | P Value |
| **Demographics** |  |  |  |  |  |  |
| Age | 34.23 (10.8) | 33.8 (10.5) | 0.83 | 36.76 (11.1) | 31.79 (10.3) | 0.062 |
| Male sex, No. (%) | 17 (20) | 12 (27) | 0.37 | 7 (24) | 6 (16) | 0.39 |
| Site Zürich, No. (%) | 61 (73) | 27 (61) | 0.19 | 20 (69) | 28 (74) | 0.67 |
| **Clinical measures** |  |  |  |  |  |  |
| Number of prior episodes | - | - | - | 2.76 (1.8) | 2.32 (1.6) | 0.29 |
| Residual depression<sup>b</sup> | 3.38 (3.7) | 0.70 (1.1) | <0.001 | 3.52 (5.14) | 2.87 (2.3) | 0.49 |
| Disease severity<sup>c</sup> | - | - | - | 0.08 (0.4) | -0.05 (0.33) | 0.14 |
| Medication load<sup>c</sup> | - | - | - | 0.007 (0.004) | 0.008 (0.004) | 0.33 |
| Month of ADM intake<sup>d</sup> | - | - | - | 24 | 28 | 0.77 |
| Psychotherapy<sup>c</sup> | - | - | - | 0.40 (0.39) | 0.39 (0.39) | 1 |
| Comorbidities<sup>b</sup> | - | - | - | 0.72 (1.03) | 0.61 (1.03) | 0.64 |
| Days of tapering | - | - | - | 52.86 (40.18) | 51.53 (43.74) | 0.90 |
| **Covariates of interest** |  |  |  |  |  |  |
| Brooding<sup>b</sup> | 9.65 (2.6) | 8.14 (2.3) | 0.001 | 10.55 (2.77) | 8.71 (1.9) | 0.002 |
| **Imaging measures** |  |  |  |  |  |  |
| Framewise displacement | 0.11 (0.05) | 0.10 (0.05) | 0.42 | 0.12 (0.07) | 0.11 (0.04) | 0.37 |

a) Unless stated otherwise, mean (SD) are shown; b) Determined as follows: residual depression: Inventory of Depressive Symptomatology-Clinician Rated(45); brooding: brooding subscale of the German version of the Response Style Questionnaire(46); comorbidities: number of past and present psychiatric diagnoses; c) Computation of the variables is described in the supplementary section S1.3; d) Median is provided and a non-parametric Wilcoxon rank sum test was applied to test significance.

### 3.2 Motion and site effects

Comparing participants with high and low motion during MA1 yielded no significant difference between these groups for any of the seeds. Hence, we decided to only exclude participants with FD > 1 between two volumes but added no additional motion regressors and did not censor any scans with stick regressors.

There was an effect of site on the temporal signal-to-noise ratio (t(126)=7.76, p<0.001, Figure S3). To examine if this impacted significant effects, we correlate the significant RSFC with site as covariate of no interest.

### 3.3 Overall functional connectivity patterns

As a sanity check we computed the connectivity of all left seeds (left sgACC, left PCC, left dlPFC and left amygdala) with all voxel in the affective mask (including voxels from the DMN, the CEN, the SN, the amygdala and the hippocampus) across all participants at MA1. The connectivity pattern replicated the patterns established in the literature (Figure S4).

### 3.4 The remitted and medicated state

To identify abnormal RSFC in the remitted medicated state, we compared remitted patients on ADM and controls. For this analysis, we focused on connectivity between the left sgACC, i.e. the main a-priori seed, with voxels in the affective mask. RSFC between the seed with voxels in the affective mask did not differ between remitted patients on ADM and controls. No differences were found for any of the exploratory seeds (right sgACC, left PCC, right and left dlPFC and right and left amygdala).

### 3.5 Relapse effects

The left sgACC was again designated as the main a-priori seed to examine associations of RSFC with relapse. RSFC between the left sgACC and voxels in the affective mask at MA1 did not differ between patients who did and did not go on to relapse during the follow-up period. No differences were found for any of the exploratory seeds.

### 3.6 Discontinuation effects

The left PCC was designated as the main a-priori seed to examine effects of discontinuation. RSFC of the left PCC with voxels in the affective mask did not show an interaction between time point of assessment (MA1 or MA2) and discontinuation group (MA1-D-MA2 and MA1-MA2-D). No interaction effects were found for any of the exploratory seeds.

### 3.7 Interaction of discontinuation and relapse

To examine the interaction between changes in RSFC before and after discontinuation and subsequent relapse, we first focused on the changes in the connectivity of the left PCC. These analysis was conducted in group MA1-D-MA2 only. No changes were identified for RSFC of the left PCC. However, examining the exploratory seeds a significant interaction between discontinuation and relapse emerged for RSFC between the right dlPFC and two peaks in the right posterior DMN, namely in the parietal cortex (Figure 3A,B; Table 2) and the posterior cingulate cortex (Figure 3A,D). Post-hoc tests did not reveal differences between relapsers and non-relapsers before discontinuation. The significant interaction was driven by an increase in RSFC in patients who discontinued but did not go on to relapse: RSFC between the right dlPFC and the right parietal cortex increased with discontinuation in those who remained well (Figure 3A,C; note the slight shift in the parietal cortex for the post-hoc test depicted in blue), while it decreased numerically without reaching statistical significance in those who went on to relapse. None of the post-hoc tests on the connectivity between right dlPFC and PCC reached significance, although a similar pattern of results is visible as for the connectivity of right dlPFC and parietal cortex (compare Figure 3B and D). The effects did not remain significant when including participants with more than 1mm FD and censuring the scans with stick regressors.

**Figure 3.**
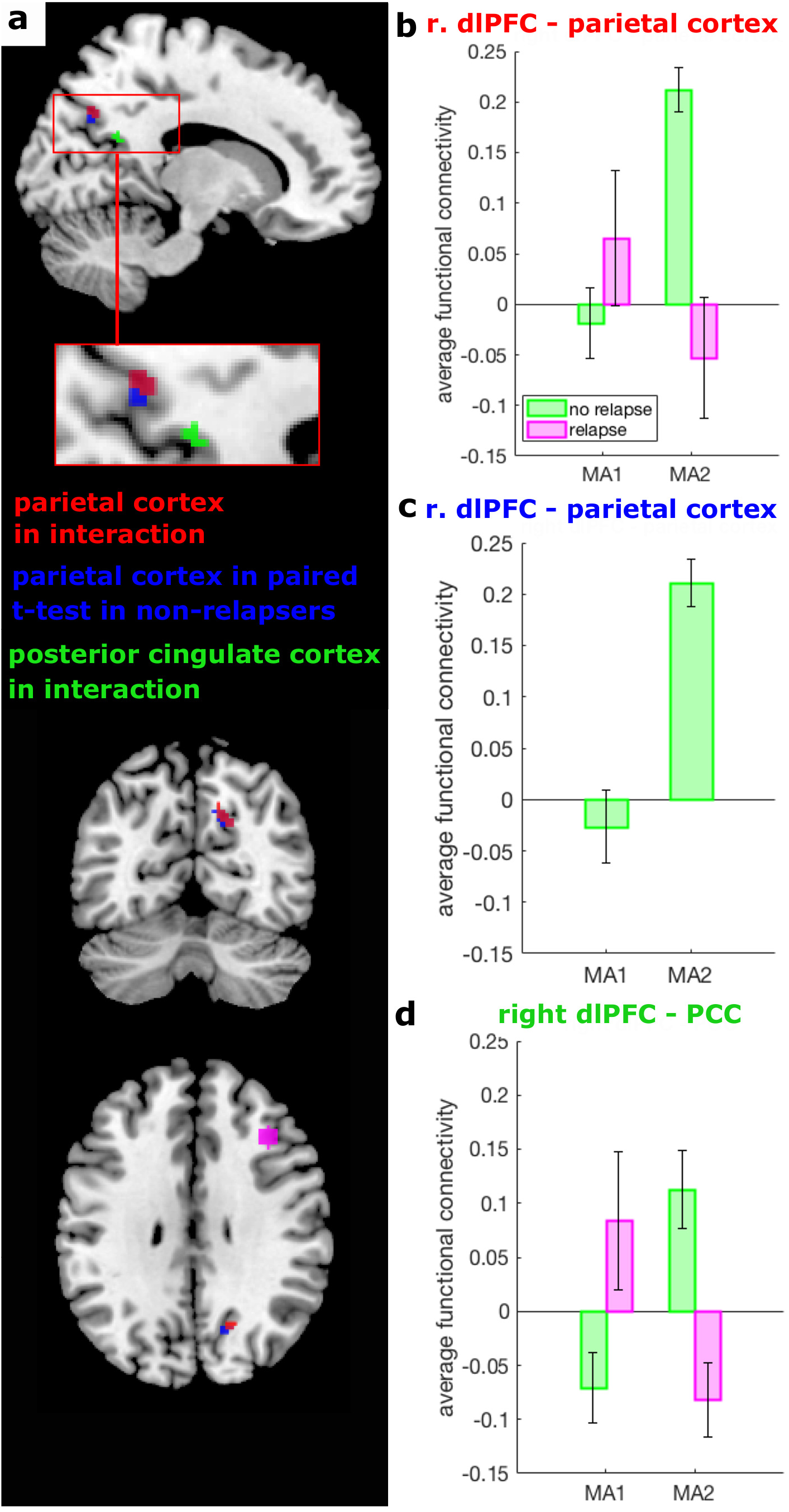
Discontinuation relapse interaction effect: A) Depicted are clusters for significant interactions between discontinuation and relapse between ourchoosen seed in the right dorsolateral prefrontal cortex (dlPFC, magenta) and the parietal cortex (red) and the posterior cingulate cortex (PCC, green) and a paired t-test in non-relapsers only for RSFC between right dlPFC and the parietal cortex (blue). B-D) The Fisher z-transformed average functional connectivity between the right dlPFC and the surviving clusters in the parietal cortex and the PCC is depicted for the interaction between relapsers and non-relapsers (B and D, respectively). Specifically, depicted are significant interaction effects with p<0.0083 which survive our threshold corrected for multiple comparison. Exact p-values are given in Table 2. C) The paired post-hoc t-test in non-relapsers for the interaction effect in relation to the parietal cortex shown in (B). Error bars indicate standard errors.

**Table 2:** Significant discontinuation relapse effects for the dorsolateral prefrontal cortex

| Contrast | Region (BA) | Peak MNI coordinates |  |  | k | p<br>(cluster) | T-Value | Z | p<br>(peak) |
| --- | --- | --- | --- | --- | --- | --- | --- | --- | --- |
|  |  | x | y | z |  |  |  |  |  |
| Interaction in F-test | Parietal cortex (BA 7) | 14 | -66 | 38 | 30 | 0.002 | 5.05 | 5.21 | 0.001 |
|  | PCC (BA 23) | 12 | -54 | 14 | 18 | 0.006 | 5.06 | 5.2 | 0.001 |
| Paired T-test<br>in non-relapsers | Parietal cortex (BA 7) | 14 | -66 | 36 | 30 | < 0.001 | 8.2 | 5.43 | 0.001 |

BA = Brodmann area, MNI = Montreal Neurological Institute, PCC = posterior cingulate cortex

### 3.8 Post-hoc prediction analyses

Using the average functional connectivity between the right dlPFC and the cluster in the parietal cortex (shown in red in Figure 3A) at MA2, relapse could be predicted out-of-sample with a balanced accuracy of 0.86 (Figure S5). Similarly, using the change in that RSFC from MA1 to MA2 also predicted relapse with a balanced accuracy of 0.86. Note that the data used in the second prediction analysis is not orthogonal to the significant interaction effect and, hence, does not provide independent evidence for the prediction.

### 3.9 Exploratory analyses without correction for multiple comparisons

No significant effects emerged for any of the exploratory seeds in any of the analyses when we dropped the correction for multiple comparison for number of seeds and set the required significance level to 0.05 FWE-corrected. In Table S1, we additionally report results for RSFC for our main seeds for all analyses at an uncorrected significance level of 0.001.

### 3.10 Covariates

We correlated RSFC between the right dlPFC and the cluster in the parietal cortex in non-relapsers before they discontinued as well as the change accompanying the discontinuation with our pre-specified covariates, but found no significant correlations.

## 4 Discussion

We examined resting-state functional connectivity in the context of antidepressant discontinuation in stable remission from Major Depressive Disorder. We found a significant interaction of discontinuation and relapse in the subgroup of patients who were tested before and after discontinuation. This interaction was driven by those who remained well during the follow-up period showing a significant increase in RSFC between the dlPFC and two areas in the posterior DMN, namely the parietal cortex and the PCC, whereas patients who went on to relapse did not show this increase and instead showed a numerical decrease that did not reach statistical significance. Both dlPFC-parietal cortex RSFC after discontinuation and the change in RSFC were predictive of relapse in leave-one-out cross-validation analyses.

There are also several negative results. First, we found no difference between remitted, medicated patients and matched controls. Second, patients who went on to relapse after ADM discontinuation did not differ from those who remained well. Third, discontinuation did not result in changes in RSFC between the PCC and voxels within our affective mask, nor any of the exploratory seeds.

Our data suggests that an increase in RSFC between the right dlPFC and the posterior DMN, in particular the parietal cortex, relates to resilience to relapse after ADM discontinuation. An effect of this kind was expected a priori as it reflects prior findings showing a decrease in PCC connectivity with ADM initiation(36; 31; 33). However, unlike previous reports of an increase in the RSFC of the CEN in those who go on to relapse(27), we here observed an increase in right dlPFC-parietal connectivity in patients who remained well. These findings hence point towards a compensatory effect in response to discontinuation that might protect patients from relapse. While the default mode has been related to rumination, the right dlPFC has been extensively related to regulation(47) and conceptual-emotional information integration(48). It is hence tempting to speculate that the increase in connectivity might reflect emotion regulation strategies and rumination, both of which influence the course of depression(49; 50).

Exploratory post-hoc analyses indicated that RSFC between the right dlPFC and the significant cluster in the parietal cortex can predict relapse risk above chance for new, previously unseen patients. The analysis suggests that the connectivity effect may not only be associated with relapse, but may also have some predictive potential. In fact, the measure of RSFC after discontinuation is sufficient to predict subsequent relapse, indicating that one scanning session may be sufficient. Being able to predict relapse risk for new patients is crucial for translation to the clinical setting, where clinicians need to make treatment decisions using models that were developed without the patient whose treatment course they need to decide upon. However, the present result also needs to be interpreted with great caution. First, the analysis was exploratory and focused on significant clusters. Second, the sample size included in this analysis was very small, which can inflate the result(51). Hence, the result can be seen as a first hint at a potential predictive marker of relapse risk which, however, requires replication in an independent and larger dataset.

We also emphasize the difficulties inherent in interpreting the absence of an effect, and we note the problems around reliability of RSFC measures for brief scans as in our study(52). Although these results do not provide evidence for the absence of effects, power considerations can nevertheless inform the interpretation by indicating the size of effects a study of this size should have identified. Two-tailed two-sample t-tests comparing 84 patients to 44 healthy controls and 29 relapsers to 38 non-relapsers have a power of 0.8 to detect medium (d=0.52) and large (d=0.70) group differences, respectively, and a power of 0.95 to detect effects of d=0.67 and 0.90, respectively (using G*Power 3.1(53)). As such, large effects above 0.5 are unlikely, and hence we judge it unlikely that resting-state connectivity revealed with a seed-based approach based on short scans as in this study have clinical utility(54).

The lack of replications of previous results could relate to specific difference in the sample or due to methodological limitations. At the present stage, it is not possible to disentangle these aspects. Hence, we will briefly point out difference in relation to the sample and discuss methodological limitations in the next section.

The absence of significant RSFC differences between remitted MDD patients on ADM and healthy never-depressed controls contrasts with results in the literature(19; 24). The only major difference between previous and the present study is the medication status: all previous involving remitted patients focused on unmedicated samples(15; 17; 19; 20; 21; 24), while our patient group had been medicated for a median duration of two years.

The absence of differences in RSFC between patients who went on to relapse and patients who remained well after ADM discontinuation is similarly surprising, given that relapse and resilience independent of discontinuation have been reported to be associated with abnormal RSFC of the sgACC(27; 26). We again note that our sample was assessed in the medicated state before discontinuation. The absence of a difference in our study raises the possibility that previous findings relate more broadly to relapse risk, and that our lack of findings speaks only to the specific induction of relapse through discontinuation.

Next, ADM discontinuation did not reveal any effects. This result, again, is unexpected given that ADM initiation has been shown to induce changes in the connectivity of the PCC(36; 31; 33). One potential explanation could be that the timing of the second measurement five half-lives after ADM discontinuation was too early to see effects.

This would in turn suggest a normalisation of RSFC lasting into the unmedicated state.

## 4.1 Limitations and Strengths

The study has both strength and limitations. Amongst its strengths is that, to our knowledge, this is the first study examining neurobiological effects in the context of ADM discontinuation and relapse thereafter. The lack of findings and replications in our study, however, might result from some of its limitations. First, the restingstate scan was only 5.5min, which is relatively short. Second, the evidence in favour of abnormal RSFC of the sgACC, on which we based our hypotheses and analyses, is admittedly heterogeneous and also does rely on partially overlapping samples(15; 19; 20; 21). Third, the signal-to-noise ratio differed between the study sites, potentially reducing the power to find effects in the bigger sample from Zurich. Fourth, the fact that patients were on medication from several classes might have increased the heterogeneity of the medications’ influence on RSFC and in that it reduced the power to identify significant effects. Fifth, no standard approach for motion correction exists. Hence, cut-off choices are arbitrary. To avoid spurious correlation due to motion(55; 56), we choose a priori a more conservative criterion by excluding all participants with more than 1mm FD. Sixth, the naturalistic design does not allow us to disentangle pharmacological effects of discontinuation from the potential psychological effect of knowing that the ADM is discontinued. Seventh, the sample size in particular for relapse effect was rather small. Nevertheless, as discussed above, it should be sufficient to detect moderate effects, which we did not find. Finally, these analyses followed previous studies of RSFC in depression, which primarily used a seed-based approach. More advanced analyses could yet reveal meaningful changes.

## 4.2 Conclusion

We found no RSFC differences between remitted, medicated patients and healthy controls, between relapsers and non-relapsers or as a consequence of discontinuation. This raises the possibility - but emphatically does not show - that antidepressants normalise RSFC and that this normalisation persists at least for a brief period into the unmedicated state. We did, however, find tentative evidence for a potential marker of resilience to relapse after ADM discontinuation in terms of an increased connectivity between the right dlPFC and the posterior DMN.

## Acknowledgements

This study was principally funded by a Swiss National Science Foundation grant (320030L_153449 / 1) to QJMH and by a Deutsche Forschungsgemeinschaft grant (WA 1539/5-1) to HW. Additional funds were provided by a the Clinical Research Priority Program “Molecular Imaging” at the University of Zurich to KES and ES. KES was additionally supported by the René and Susanne Braginsky Foundation.

## Conflicts of interest

All authors declare no conflicts of interest. The funders had no role in the design, conduct or analysis of the study and had no influence over the decision to publish.

## Contributions

QJMH and HW conceived and designed the study with critical input by KES and ES. IMB, JW, IS and LKu collected the data under the supervision of QJMH and HW. KES was the study sponsor in Zurich. QJMH, ES, KES and HW acquired funding for the study. IMB planned and performed the analyses and wrote the manuscript under supervision of QJMH. LKa and IMV gave advice on the analyses. All authors provided critical comments, read and approved the manuscript.

## Data Availability

The datasets generated and analysed during the current study are not publicly available due to constrictions by the ethical rules to but are available from the corresponding author on reasonable request and in line with the ethical rules.

## Supplementary Material

## S1 Supplementary Methods

### S1.1 In- and Exclusion Criteria

Participants fulfilling the following inclusion criteria were eligible for participation in the study:

1. age 18-55 years
2. ability to consent and adhere to the study protocol
3. written informed consent
4. fluent in written and spoken German.

Patients had to additionally fulfil the following criteria:

1. currently under medical care with a psychiatrist or general practitioner for remitted Major Depressive Disorder and willing to remain in care for the duration of the study (approx. 9 months)
2. informed choice to discontinue medication (including willingness to taper the medication over at most 12 weeks) that was independent of study participation
3. clinical remission (Hamilton Depression Score of less than 7) had been achieved under therapy with Antidepressant Medication (ADM) without having undergone manualized psychotherapy; with no other concurrent psychotropic medication and had been maintained for a minimum of 30 days,
4. consent to information exchange between treating physician and study team members regarding inclusion/exclusion criteria and past medical history.

Any of the following exclusion criteria led to exclusion of an participant. This included the following general criteria

1. any disease of type and severity sufficient to influence the planned measurement or to interfere with the parameters of interest (This includes neurological, endocrinological, oncological comorbidities, a history of traumatic or other brain injury, neurosurgery or longer loss of consciousness.)
2. premenstrual syndrome (ICD-10 N94.3).

and MRI-related criteria

1. MRI-incompatible metal parts in the body,
2. inability to sit or lie still for a longer period,
3. possibility of presence of any metal fragments in the body,
4. pregnancy,
5. pacemaker, neurostimulator or any other head or heart implants,
6. claustrophobia and
7. dependence on hearing aid.

For patients the following additional criteria would led to exclusion:

1. current psychotropic medication other than antidepressants,
2. questionable history of major depressive episodes without complicating factors,
3. current acute suicidality,
4. lifetime or current axis II diagnosis of borderline or antisocial personality disorder,
5. lifetime or current psychotic disorder of any kind, bipolar disorder,
6. current posttraumatic stress disorder, obsessive compulsive disorder, or eating disorder
7. current drug use disorder (with the exception of nicotine) or within the past 5 years.

Healthy controls were excluded if there was a lifetime history of Diagnostic and Statistical Manual of Mental Disorders (4th ed., text rev.)(1) axis I or axis II disorder with the exception of nicotine dependence.

### S1.2 Questionnaires and Clinical Assessments

Clinical in- and exclusion criteria were assessed with the Structured Clinical Interview for DSM-IV (SCID) I and II to diagnose axis 1 disorders (major mental disorders) and axis II disorders (personality disorders), respectively(2). The Structured Interview Guide for Hamilton Depression Rating Scale (SIGH-D)(3) consisting of 17 items was used to assess inclusion and the Inventory of Depressive Symptomatology Clinician Rated (IDS-C)(4) with 30 items to quantify residual depression. Additionally, we applied the German version of the Response Style Questionnaire (RSQ-10D)(5) measuring brooding and reflection as components of rumination with 5 items each.

### S1.3 Data Analysis

All analyses, except for the preprocessing of the imaging data, were performed using Matlab version 2016b.

We computed an overall measure of disease severity as the first principal component of number of past depressive episodes, age at illness onset, time in remission, time since depression onset, severity of last episode, time sick in total and time sick in the last five years as variables.

Medication load was based on the dose prior to discontinuation divided by the maximal allowed dose according to the Swiss compendium (www.compendium.ch) and by the weight of the participant.

Psychotherapy score was coded such that patients with no psychotherapy within the year before the study received a 0, patients reporting to have completed a psychotherapy within one year before the study a 0.5 and patients reporting to be in psychotherapy at the beginning of the study as 1. Significance was computed with a three-way chi-squared test.

### S1.4 Image Acquisition

Images were acquired at the two study sites using a Phillips 3T Ingenia in Zurich and a Siemens 3T Trio in Berlin. Participants were instructed to stay awake, keep their eyes open and look at a centrally placed fixation cross.

In Zurich, a 32-channel coil was used to acquire echo-planar images (EPIs; 136 volumes; 40 axial slices; 2.5mm slice sickness; descending sequential acquisition, repetition time: 2560 ms; echo time: 27, field of view: 210 x 210 x 119.5, acquisition matrix: 84 x 82, reconstructed voxel size: 2.19 x 2.19 x 2.50 mm, flip angle: 90°). Additionally, we acquired T1-weighted magnetization-prepared rapid-acquisition gradient-echo (MPRAGE) structural images (301 axial slices; slice thickness: 1; repetition time: 7.9ms; echo time: 3.7ms, field of view: 250 x 250 x 180.6, acquisition matrix: 252 x 251, reconstructed voxel size: 0.98 x 0.98 x 0.60 mm, flip angle: 8^°^).

In Berlin, a 32-channel coil was used for functional resting-state EPIs (136 volumes; 40 axial slices; 3mm slice thickness including a gap of 0.5mm; descending sequential acquisition, repetition time: 2560 ms; echo time: 27 ms, field of view: 210 x 210 x 120, acquisition matrix: 84 x 84, voxel size: 2.50 x 2.50 x 2.50 mm, flip angle: 90°). T1-weighted MPRAGE structural images (192 axial slices; slice thickness: 1mm; repetition time: 1900 ms; echo time: 2.52 ms, field of view: 256 x 256 x 192, acquisition matrix: 256 x 256, reconstructed voxel size: 0.98 x 0.98 x 0.60mm, flip angle: 9°) were also acquired.

### S1.5 Preprocessing

Functional images were realigned, slice-time corrected and smoothed with a 6mm FWHM kernel using adaptive spatial procedure (SUSAN(6)) in FSL (FMRIB Software Library v5.0). The images were then co-registered to the structural image and normalised using Advanced Normalization Tools (ANTs(7)). Finally, an independent component analysis-based artefact removal (ICA-AROMA(8)) was applied to exclude noise components relating e.g. to breathing and heart rate, using a data-driven approach and the data was subjected to a high-pass filter of 0.008Hz. Lastly, BOLD data were normalised to MNI standard space, applying the registration matrices and warp images from the two previous registration steps, and then resampled into 2 mm isotropic voxels. All imaging data were visually inspected to exclude acquisition artefacts or other data corruption.

### S1.6 Motion correction

As group differences can be confounded by head motion differences(11), we excluded participants from all analyses if their frame-wise displacement (FD) from one volume to the next exceeded 1mm at any time during the scan. To test for the effects of motion, we performed a median-split based on the mean FD and compared RSFC for all seeds between all participants included at MA1. In case effects were negligible, we used 6 realignment parameters as motion regressors on the first level and no further correction to avoid over-fitting and power reduction. In case non-negligible motion artefacts were observed, we would have additionally added the 6 derivatives of the realignment parameters and censored those scans for which FDs were bigger than 0.5. Censoring scans means to include an additional regressor for each volume at which the movement exceeds a given threshold, here 0.5 FD. This regressors contains zeros at all volumes but the volume that exceeds the threshold. At that volume, the regressor contains a one.

### S1.7 Study site effects

To examine systematic differences between the two study sites, we compared the temporal signal-to-noise ratio in the grey matter for all included subjects between sites.

### S1.8 Affective mask creation

The “affective mask” consists of functional and anatomical masks that were merged in SPM.

The following regions of interest (ROIs) were taken from the CONN toolbox(9) to build masks for the default mode network, the salience network and the executive (dorsal attention and fronto parietal) network defined by ICA analyses of 497 subjects from the human connectome project in the toolbox.

DefaultMode.MPFC (1,55,-3)

DefaultMode.LP (L) (−39,−77,33)

DefaultMode.LP (R) (47,−67,29)

DefaultMode.PCC (1,−61,38)

Salience.ACC (0,22,35)

Salience.AInsula (L) (−44,13,1)

Salience.AInsula (R) (47,14,0)

Salience.RPFC (L) (−32,45,27)

Salience.RPFC (R) (32,46,27)

Salience.SMG (L) (−60,−39,31)

Salience.SMG (R) (62,−35,32)

DorsalAttention.FEF (L) (−27,−9,64)

DorsalAttention.FEF (R) (30,−6,64)

DorsalAttention.IPS (L) (−39,−43,52)

DorsalAttention.IPS (R) (39,−42,54)

FrontoParietal.LPFC (L) (−43,33,28)

FrontoParietal.PPC (L) (−46,−58,49)

FrontoParietal.LPFC (R) (41,38,30)

FrontoParietal.PPC (R) (52,−52,45)

Amygdala and hippocampus masks:

These masks were built using the SPM Anatomy toolbox(10). We used anatomical ROIs to create the right and left amygdala labelled AStr, CM, LB and SF in the toolbox. Similarly, we used anatomical ROIs to create the right and left hippocampus labelled CA1, CA2, CA3 and DG in the toolbox.

### S1.9 Sanity checks and exploratory analyses

To specifically examine effects of time, paired t-test in patients who did not discontinue but were assessed twice (group MA1-MA2-D) were conducted.

To ensure the validity of our method, we repeated the analyses without adding the covariates from aCompCor in the first level. We also repeated the analyses without adding motion regressors at that stage.

To explore whether we missed strong abnormalities that were outside our restricted search volume, i.e. the affective mask, which might be of interest for future studies, we repeated all second level analyses without the affective mask in whole-brain analyses. In addition, we report results without correction for multiple comparison for number of seeds and uncorrected results at a significance level of 0.001 for all main seed analyses to allow for estimates of potential type II errors.

## S2 Supplementary Results

### S2.1 Quality checks

To ensure that functional ectivity between our chosen seeds and the anticipated networks based on the literature was evident, we visually inspected the networks connected to the seeds in all participants included for analyses

**Figure S1.**
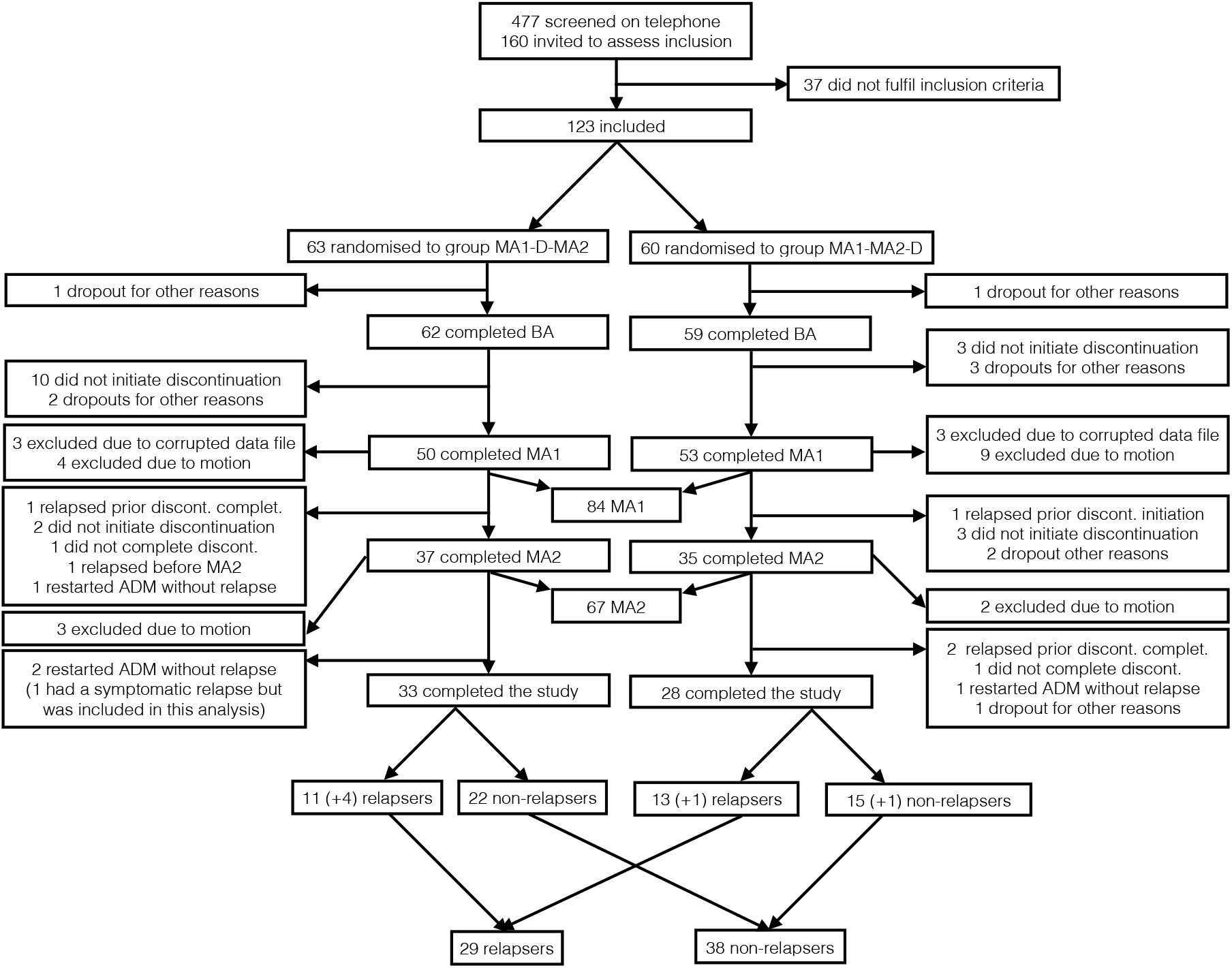
Consort Diagram for Patients: Depicted are reasons for dropouts and exclusion for patients throughout the study. (+ X) indicates the number of participants who either relapsed or did not relapse but did not have useable data at main assessment (MA) 2. MA1-D-MA2 = Discontinuation between MA1 and MA2; MA1-MA2-D = MA1 and MA2 before Discontinuation; BA = baseline assessment

**Figure S2.**
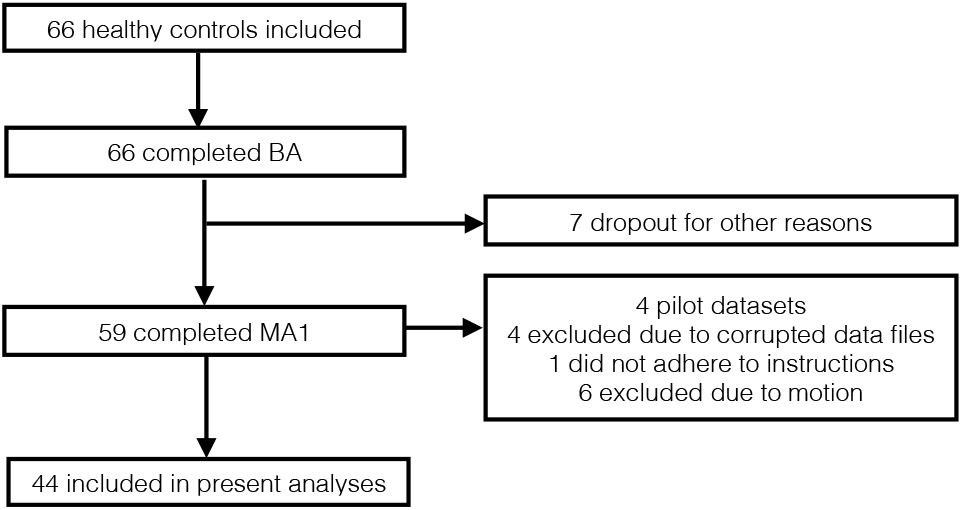
Consort Diagram for Healthy Controls: Depicted are reasons for dropouts and exclusion for controls throughout the study. BA = baseline assessment; MA = main assessment

**Figure S3.**
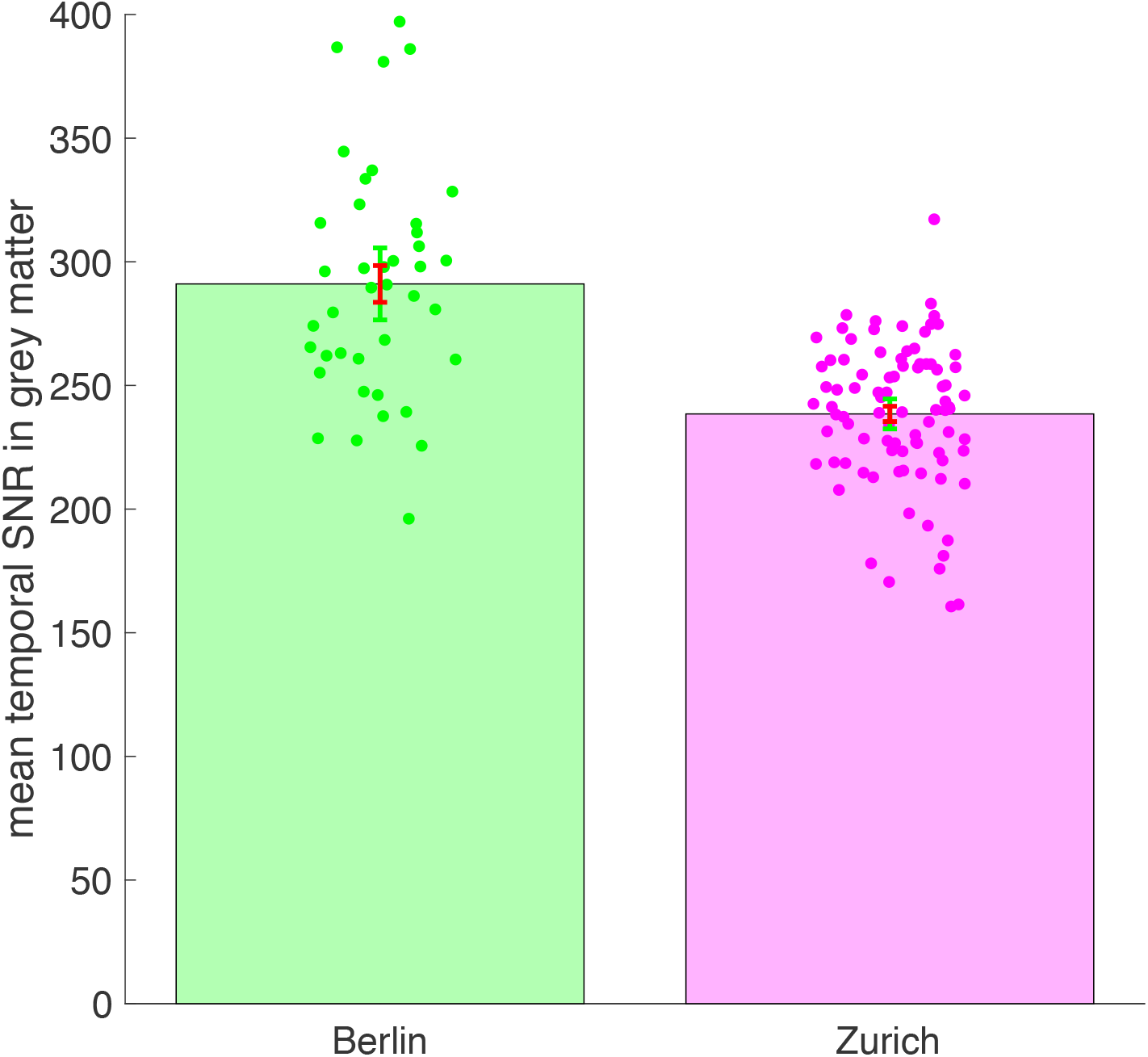
Site effects: Depicted is the average signal-to-noise ratio (SNR) within the individual grey matter masks over the time of the resting-state period. Dots indicate individual data points, red error bars show standard errors and green error bars show 95% confidence intervals.

at MA1. Figure S4 depicts these networks for all seeds in the left hemisphere. Network functional connectivity seems as expected.

**Figure S4.**
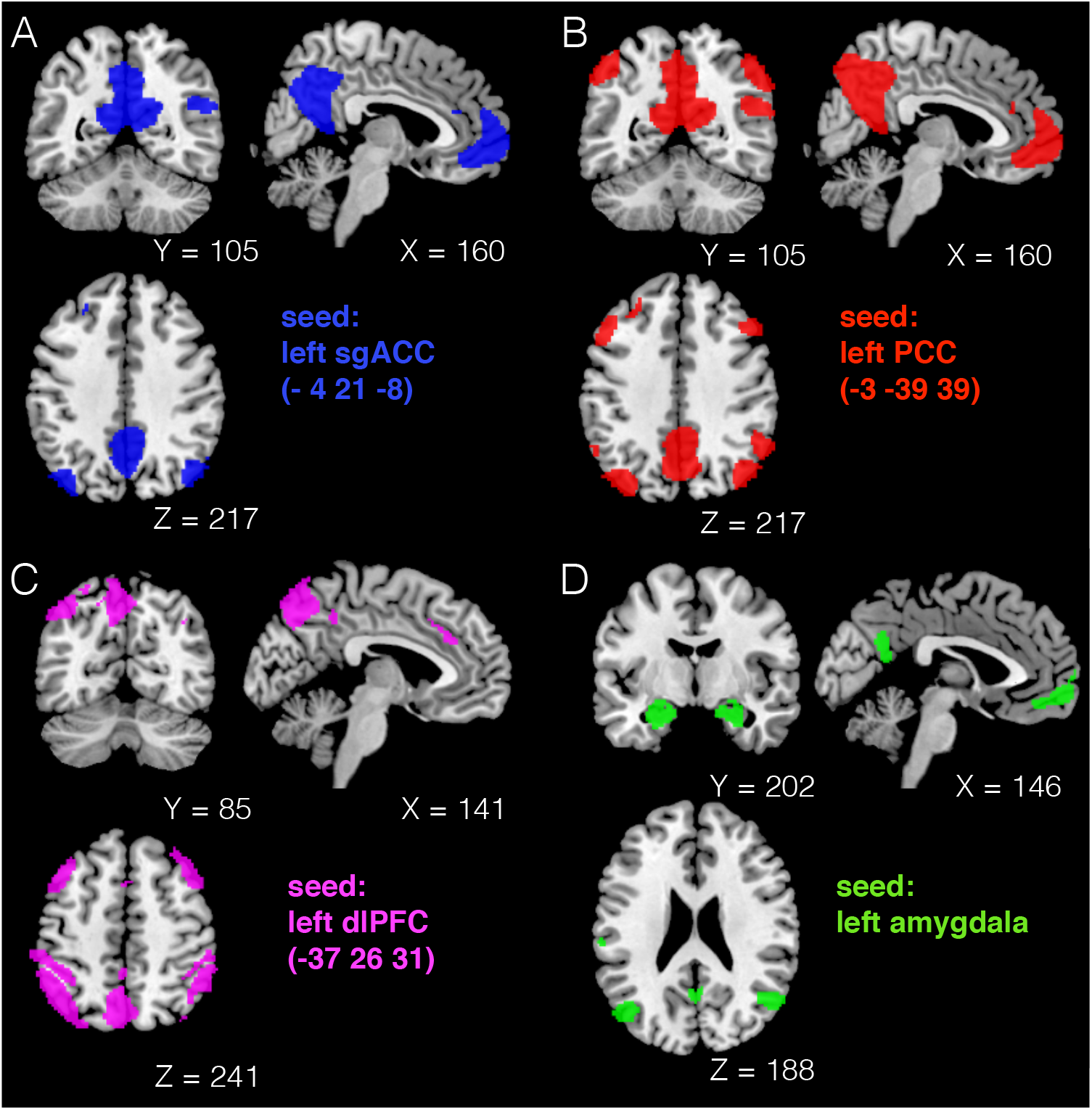
Functional connectivity networks of all left-sided seeds: sgACC = subgenual anterior cingulate cortex; PCC = posterior cingulate cortex, dlPFC = dorsolateral prefrontal cortex

### S2.2 Effects of time

There were no significant changes in RSFC for any of the seeds in patients who were assessed twice prior to discontinuation.

### S2.3 Effects of noise regressors on the first level

Analyses without regressors for motion on the first level replicated the main pattern of results. Not including additional regressors from aCompCor in the first level analyses also replicated the main pattern of results.

### S2.4 Whole-brain exploratory analyses

Repeating all second level analyses without the affective mask led to the similar significant clusters as reported for the within mask analyses, whereas the p-values naturally differed (parietal cortex: p=0.021, PCC: p=0.004). Of note, no additional effects emerged.

### S2.5 Uncorrected results

Table S1 depicts results for all main seeds considered significant at 0.001 without correction. The sparsity of results at this significance level speaks against a high rate of type II error due to correction for multiple comparison, but supports the null hypotheses for many of the examined effects.

**Table S1:** Significant results at uncorrected level for main seeds

| Ssed | Contrast | BA/Region | Peak MNI coordinates |  |  | k | p<br>(uncor.) | T-Value | Z | p<br>(FWE-cor., peak) |
| --- | --- | --- | --- | --- | --- | --- | --- | --- | --- | --- |
|  |  | x | y | z |  |  |  |  |  |  |
| left sgACC | T-test for Pat - HC | Left BA 10 | -34 | 48 | 14 | 1 | 0.001 | 3.26 | 3.19 | 0.821 |
|  | T-test for HC - Pat | Left BA 8 | -34 | 22 | 58 | 1 | 0.001 | 3.30 | 3.22 | 0.795 |
|  | T-test for No Rel - Rel | Right BA 19 | 48 | -72 | 26 | 11 | < 0.001 | 3.60 | 3.45 | 0.612 |
|  | T-test for No Rel - Rel | Right sensory assoc. | 28 | -42 | 64 | 1 | 0.001 | 3.25 | 3.11 | 0.942 |
| left PCC |  |  |  |  |  |  |  |  |  |  |
|  | F-test discontinuation | Left BA 7 | -38 | -48 | 56 | 26 | < 0.001 | 4.35 | 4.19 | 0.059 |
|  | F-test discontinuation | Right amygdala | 16 | -2 | -18 | 8 | < 0.001 | 3.56 | 3.47 | 0.522 |
|  | F-test discontinuation | Right amygdala | 24 | 4 | -26 | 2 | < 0.001 | 3.53 | 3.44 | 0.556 |
|  | F-test discontin. x rel. | Right BA 10 | 40 | 58 | -8 | 3 | < 0.001 | 3.78 | 3.57 | 0.432 |

T-tests are independent sample t-tests. F-tests show interactions for the discontinuation effect and the discontinuation x relapse interaction effect. BA = Brodmann area, MNI = Montreal Neurological Institute, sgACC = subgenual anterior cingulate cortex, HC = healthy controls, Pat. = patients, No Rel = no relapse, Rel = relapse, PCC = posterior cingulate cortex

**Figure S5.**
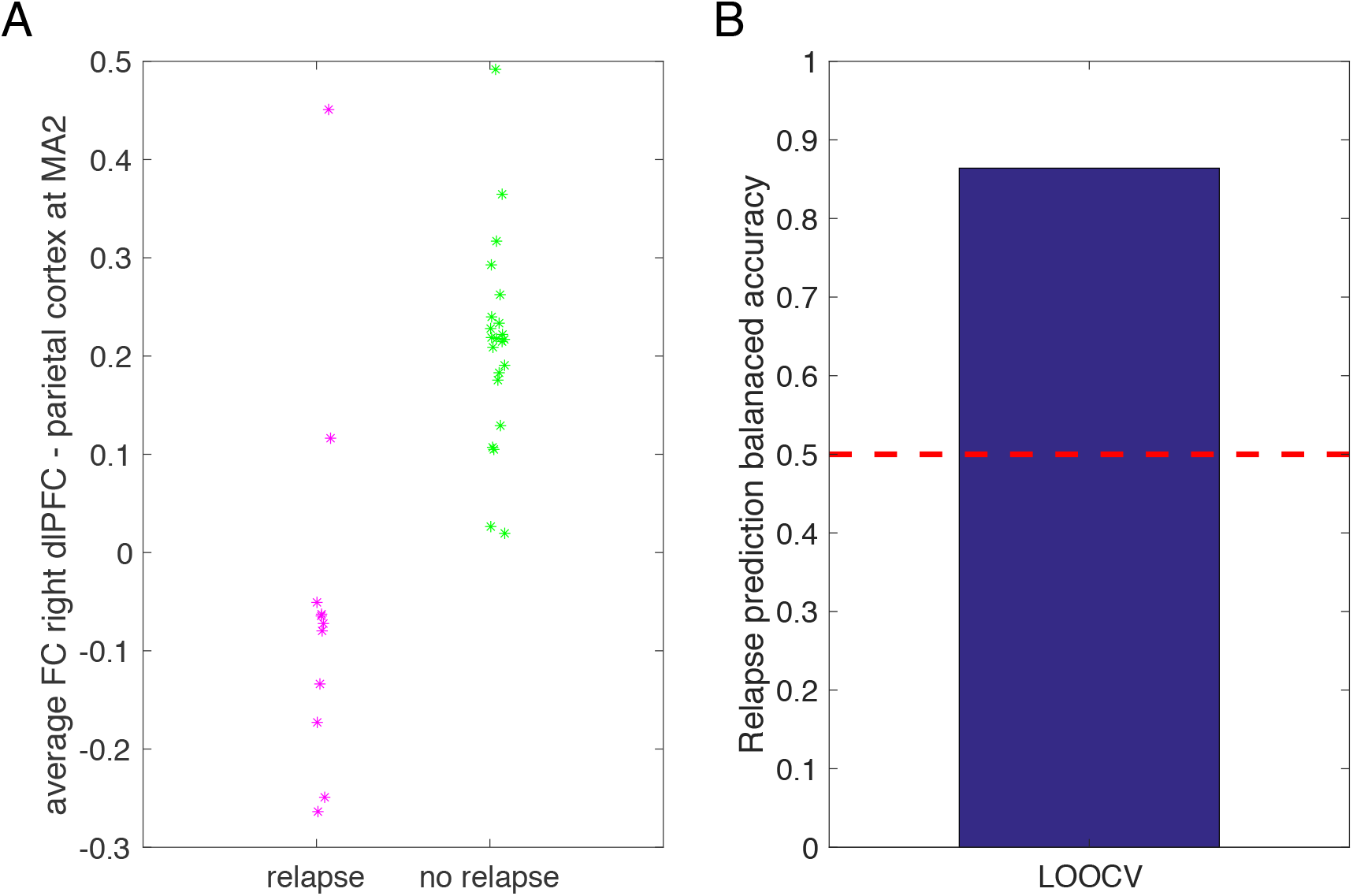
Prediction analyses: A) Average functional connectivity (FC) for the significant cluster in the parietal cortex at main assessment 2 (MA2) B) Balanced accuracy for predicting subsequent relapse using leave-one-out cross-validation. The dashed red line indicates chance level. dlPFC = dorsolateral prefrontal cortex

## Notes

### Competing Interest Statement

The authors have declared no competing interest.

